# Effect of OTR4120 on pulmonary fibrosis

**DOI:** 10.1101/2021.10.24.465650

**Authors:** Said Charef, Najat Mejdoubi Charef, Franck Chiappini, Dulce Papy-Garcia, Denis Barritault

## Abstract

**Background:** Acute respiratory distress syndrome (ARDS) can be related to airway remodeling caused by pulmonary fibrosis and systemic inflammation. Etiologies of ARDS are multifaceted such as idiopathic pulmonary fibrosis or as recently the SARS-CoV-2 infection. Antifibrotic drugs may be a better approach to slow the fibrotic process but they often have poor efficacy in patients, and the mortality rate remains high, up to 40% within 5 years of diagnosis.

Here, we tested the antifibrotic effect of a ReGeneTaring Agents named OTR4120 in a bleomycin-induced mouse model of pulmonary fibrosis.

**Methods:** Swiss mice were randomly divided into four experimental groups: saline-treated control group, an OTR4120 group, a bleomycin-induced fibrosis group without OTR4120, and a bleomycin-induced fibrosis groups with OTR4120 (intravenous injections every 3 days starting at day 11 post bleomycin I.P. injection). Lungs were compared using the lung/body weight index, and the extend of interstitial injury area was graded using histopathological assessment of haematoxylin & eosin-stained lung tissue sections. Lung tissue Collagen I and Collagen III levels, and blood cytokine levels were measured using a Collagen colorimetric kit and a Cytokine colorimetric kit, respectively.

**Results:** The group treated by OTR4120 alone were used as a control. The clincal signs in all animals resoved gradually on day 17 after bleomycin injections and 6 days after OTR4120 treatment, and disappeared almost completetly at day 24 after bleomycin injections and day 13 after OTR4120 treatment. Lung/body weight index values were significantly lower in the bleomycin-OTR4120 treated group versus the bleomycin group (7.31, 9.97 and 7.63 mg/g, p-value< 0.01; respectively). Histopathological analyses suggest that OTR4120 treatment ameliorated the increased inflammatory cell infiltration, and attenuated the reduction in interstitial thickening, associated with bleomycin-induced fibrosis. Collagen III and cytokine levels were decreased in the OTR4120 group versus the fibrotic (bleomycin only) group. OTR4120-treated animals were less affected in their behavior, did not loose weight nor appetite, and recovered overall activities within 6 days of OTR4120 treatment, while none of the vehicle-treated animals recovered to normal.

**Conclusion:** OTR4120 is a potential candidate to reduce lung fibrosis.

## Introduction

Pulmonary fibrosis is a common outcome of various of known or still unknown causes [1]. The so called idiopathic pulmonary fibrosis (IPF) is characterized by progressive dyspnea and loss of pulmonary function [2,3]. Median survival in patients with IPF is only 3–5 years [5], and disease related morbidity and mortality has risen gradually over recent years [5]. The mechanisms underlying the pathogenesis of pulmonary fibrosis often include the accumulation of inflammatory cells in the lungs, and the generation of pro-inflammatory and pro-fibrotic mediators, resulting in alveolar epithelial cell injury and fibroblast hyperplasia, and ultimately excessive deposition of extracellular collagen [6]. Corticosteroids and other immunosuppressive drugs are used in the treatment of some inflammatory pulmonary fibrosis but are inefficient in IPF. Thus, antifibrotic drugs may be a better approach to slow the fibrotic process [7]. However, they often have poor efficacy in patients, and the mortality rate remains high, up to 40% within 5 years of diagnosis [7]. Lung transplantation is an effective means of treating pulmonary fibrosis; however, its application is limited due to complications, infection, high costs, and particularly, a lack of donated organ resources [8].

The interest of new approaches for the treatment of pulmonary fibrosis has recently increased due to the novel human coronavirus which mainly affects the respiratory system. SARS-CoV-2, causes a respiratory disease characterized by cough (mostly dry), dyspnea, fatigue, and, in severe cases, pneumonia or respiratory failure (corroborated by radiographic bilateral ground-glass opacity) [9–11]. Damage to the airway tract and lungs was evident during biopsy and autopsy studies [9–11]. Diffuse alveolar damage (DAD) and airway inflammation have been reported both in humans and in nonhuman primates [9, 12]. The leading cause of mortality for SARS-CoV-2 is respiratory failure from acute respiratory distress syndrome (ARDS) [13]. ARDS can be related to airway remodeling caused by pulmonary fibrosis and systemic inflammation [14–16].

We are currently developing the ReGeneraTing Agent (RGTA) OTR4120, a carboxymethyl sulphated dextran, as a medicinal product [17–18]. These agents are biopolymers engineered and selected for their ability to mimic the biological activities of heparan sulfate (HS) which are key element of the extracellular matrix (ECM) scaffolding and natural sites for the storage and protecting of most communication peptides. Indeed, through their interaction with the HS Binding Site of the ECM structural proteins (such as collagens; elastin, laminin etc…), HS are key players to organize the ECM scaffold, and for the positioning and protection of communication peptides (growth factors, chemokines etc..) that regulate locally tissue homeostasis. Moreover, as a structural and functional analogous to naturally HS, one of the most crucial properties of RGTA is its resistance to enzymatic degradation [19]. Together these properties allow RGTAs to maintain their structure and activity even in the microenvironment of chronic of wounds, characterized by unrestrained proteolytic activity [19]. In injured tissue, HS, present at the cell surface and within the ECM and other ECM components, required for tissue healing and homeostasis, are destroyed by locally secreted proteases and glycanases. Introduced at the site of injury, RGTA can replace destroyed HS, reversing the hostile microenvironment and fostering tissue healing. We present the use of one specific RGTA, namely OTR4120 which was shown to enhance the speed and quality of healing in a bell-shaped dose-dependent manner after a single or weekly systemic administration of 1–2 mg/kg by various routes [17–18]. They also induce extracellular matrix remodeling and anti-fibrotic activity in several preclinical models and reduce scar formation in patients for aesthetic mastectomy [19].

The effects of OTR4120 on development of pulmonary fibrosis in response to bleomycin were evaluated compared with healthy control mice and a fibrotic (bleomycin-induced) control, with the aim of revealing potential new options for the treatment of pulmonary fibrosis in clinical drug therapy.

## Materials and methods

### Animals

Adult Swiss male mice (*n* = 20), weighing 18–22 g, were provided by Janvier Laboratories (Le Genest, St. Isle, France). The animals were housed in cages with a 12-h:12-h light–dark cycle at an ambient temperature of approximately 20–26°C and relative humidity of approximately 40-70% with free access to food and water. Animal facilities, were conducted according to the established guideline and approved by the Animal Care Committee of French University (APAFIS# 19838)

### Bleomycin fibrosis model and treatment protocol

Lung fibrosis was induced in pathogen-free male Swiss mice by two intraperitoneal (I.P.) injections of bleomycin (Bleomycin sulfate-BML-AP302-0010, Enzo Life Sciences), diluted in 0.9% saline (0.1 mg/kg BW), at days 0 and 2.

At day 11, after the acute inflammatory response, the treatments with OTR4120 (1.0 mg/kg) or 0.9 % saline (control; physiological serum) were initiated. The doses used and the protocol consisting of various i.p. injections have previously been described [20]. OTR4120 was dissolved in 0.9 % saline solution for injection.

Mice received an injection of OTR4120 or 0.9 % saline solution on days 11, 14, 17, 21 and 24, and were killed at day 28. The mice were divided randomly into the following experimental groups (five mice per group): a) 0.9 % saline only; b) OTR4120 (1.0 mg/kg starting at day 11) only; c) bleomycin (0.1 mg/kg), at day 0 and 0.1 mg/kg at day 2) with saline or d) with OTR4120 (1.0 mg/kg) starting at day 11 and repeated at days 14, 17, 21 and 24 (see **Figure 1**). The animals’ general behavior and clinical signs, including food intake, body weight and inactivity were monitored daily in each group for 28 days. To assess the food intake and body weight (BW), the mice were housed individually. Their food intake was measured by offering daily known weights of food, and separating and weighing any leftover food in the box at each change. Mice were weighed every 4 days to assess their BW.

**Figure 1.**
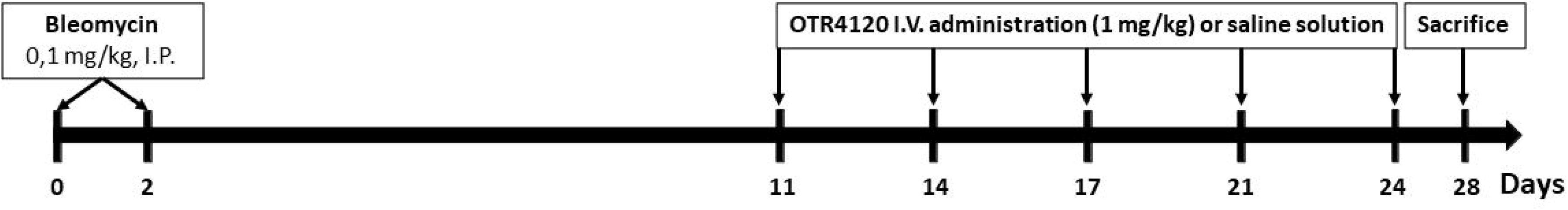
Schematic illustration of experimental design. Lung fibrosis was induced in pathogen-free male Swiss mice by two intraperitoneal (I.P.) injections of bleomycin diluted in 0.9% saline (0.1 mg/kg of BW), at days 0 and 2. At day 11, after the acute inflammatory response, the treatments with OTR4120 (1 mg/kg) or physiological serum were initiated. BW.: body weight

Pain and distress were evaluated as described by Burkholder et al. [21] with minors modifications: 0 = no signs of stress, mouse is active in good condition, calm and has normal appetite; 1 = no/mild signs of stress, mouse is active but shows some signs of restlessness; 2 = mild pain and distress, mouse is not groomed well and slightly hunched, less appetite; 3 = moderate stress, mouse moves slowly and shows signs of depression; 4 = severe pain, mouse loses significant weight and shows non-response reaction when touched, if symptoms become worse, mice were excluded from analysis and euthanized.

### Lung specimen collection

A panel of tests was used to analyze the induced pulmonary fibrosis in mice, including lung/body weight index, haematoxylin and eosin staining on lung slices (*i.e.* histopathology), Collagen I and III, TNF-α, and IL-6 assays.

After euthanasia, a skin incision was made in each mouse, along the center line of the thorax and the sternum or ribs, to expose the lungs and allow observation of the pathological lung tissue. The lung was fixed by in toto immersion in 10 % formaldehyde for 48 h, then dehydrated and paraffin-embedded using standard methods.

Tissue sections of approximately 4 μm thickness were then cut and stored at 4°C for subsequent histopathological analysis.

### Lung /body weight index

The lung/body weight index was determined according to the mass of the lungs in mg and the body weight of the mouse in g using the formula : Lung index = lung mass (mg)/body weight (g) [22].

### Histopathology

The grade of lung fibrosis was determined by histopathology, and the numerical fibrotic score was determined as described previously [23]. Briefly, the left lung was fixed, embedded, and sectioned as described above. After dewaxing with xylene, hydration was performed using a series of graded concentrations of ethanol (100% ethanol for 5 min, 95% ethanol for 1 min, 80% ethanol for 5 min, 75% ethanol for 5 min and distilled water for 2 min). Standard haematoxylin and eosin (H&E) staining was performed at room temperature for 12 min, then the sections were dehydrated and cleared using two incubations with xylene at room temperature for 10 min each. Sections sealed with neutral resin were observed under a light microscope for histopathological changes. One slide per lung was evaluated and three x20 fields of view per slide were observed. The degree of parenchymal lung remodeling was determined as previously described [24].

Alveolar septal thickening was quantified using digital imaging. At least 10 photomicrographs without overlapping across the cut surface of the lung tissue at 100 x magnification were taken for each experimental animal. Used system consists of a microscope (Axio Lab.A1, Carl Zeiss MicroImaging GmbH; Göttingen, Germany) with a video camera (AxioCam ERc5s, Carl Zeiss; Göttingen, Germany) connected to a personal computer. Gathered images were processed using the software Mesurim 2. Broncho-vascular strands were carefully removed from the analyzed areas. The software made possible to count the pixels of a given color and, knowing the area of a pixel, the area of the healthy alveoli (in percentage) could be compared to the total area of the image of the histological section.

### Bronchoalveolar lavage fluid (BALF)

The lungs were instilled by fractionated 1 ml of sterile phosphate-buffered saline (PBS) 3 times [25]. Then, the bronchoalveolar lavage fluid (BALF) was immediately centrifuged at 3000 rpm for 10 min at 4°C and stored the cell-free supernatants at −80°C for cytokine analysis. We counted the total cells, neutrophils, and macrophages with hematoxylin and eosin (H&E)-stained smears after suspending the sediment cells in PBS, and total proteins in the BALF were determined using the Pierce™ BCA Protein Assay kit according to the manufacturer’s instructions.

### Cytokine assays

We used ELISA kits to detect the concentrations of inflammatory mediators such as TNF-α (KIT ELISA TNF-a-RAB0477-1KT, Sigma-Aldrich) and IL-6 (KIT ELISA IL-6-RAB0308-1KT, Sigma-Aldrich) in BALF following the kit instructions. The optical density of each well was set at 450 nm (Tecan-Infinite, M1000 and Magellan 6 software).

### Collagen I and Collagen III assays in lung tissues

The Collagen I and Collagen III contents in lung tissue were determined by ELISA kits (Collagen, Type I (COL1) ELISA Kit-ABIN627601 and Collagen, Type III (COL3) ELISA Kit-ABIN627602 Antibodies Online). The lung tissues (100 mg) obtained were homogenized in PBS buffer (1,5 ml), centrifuged and the supernatant collected. The Collagen I and Collagen III contents in the supernatant were measured using a colorimetric assay kit (Biovision, Milpitas, CA, USA) according to the manufacturer’s instructions. Absorbance was read at 540 nm (Tecan-Infinite, M1000 and Magellan 6 software). The soluble I and III collagen levels were determined using a standard curve. Results were expressed as ng/ml collagen per lung.

### Statistical analyses

Data were presented as means ± SD. One way ANOVA was used and post-hoc t-test was used to determine the level of significance of differences between population means. Type I error was set up at 5% (*i.e.* α= 0.05). A significant difference was accepted with p-value< 0.05.

## Results

### OTR4120 reversed almost all clincal signs induced by bleomycin after 24 days

Clinical and gross pathological observations were performed: treated mice presented clinical signs over a 28-day time period after bleomycin treatment. The onset and evolution of clinical signs were as follows: on day 10 after treatment by bleomycin, slight altered gait, inactivity, inappetence. The onset of inappetence and inactivity was correlated with loss of body weight, which continued to decline (**Figure 2a** and **2b**). In the group treated by OTR4120, the clinical signs in most animals resolved gradually on days 17 after bleomycin injections and 6 days after OTR4120 treatment, and disappeared almost completely at days 24 after bleomycin injections and days 13 after OTR4120 treatment (**Figure 2c**).

**Figure 2.**
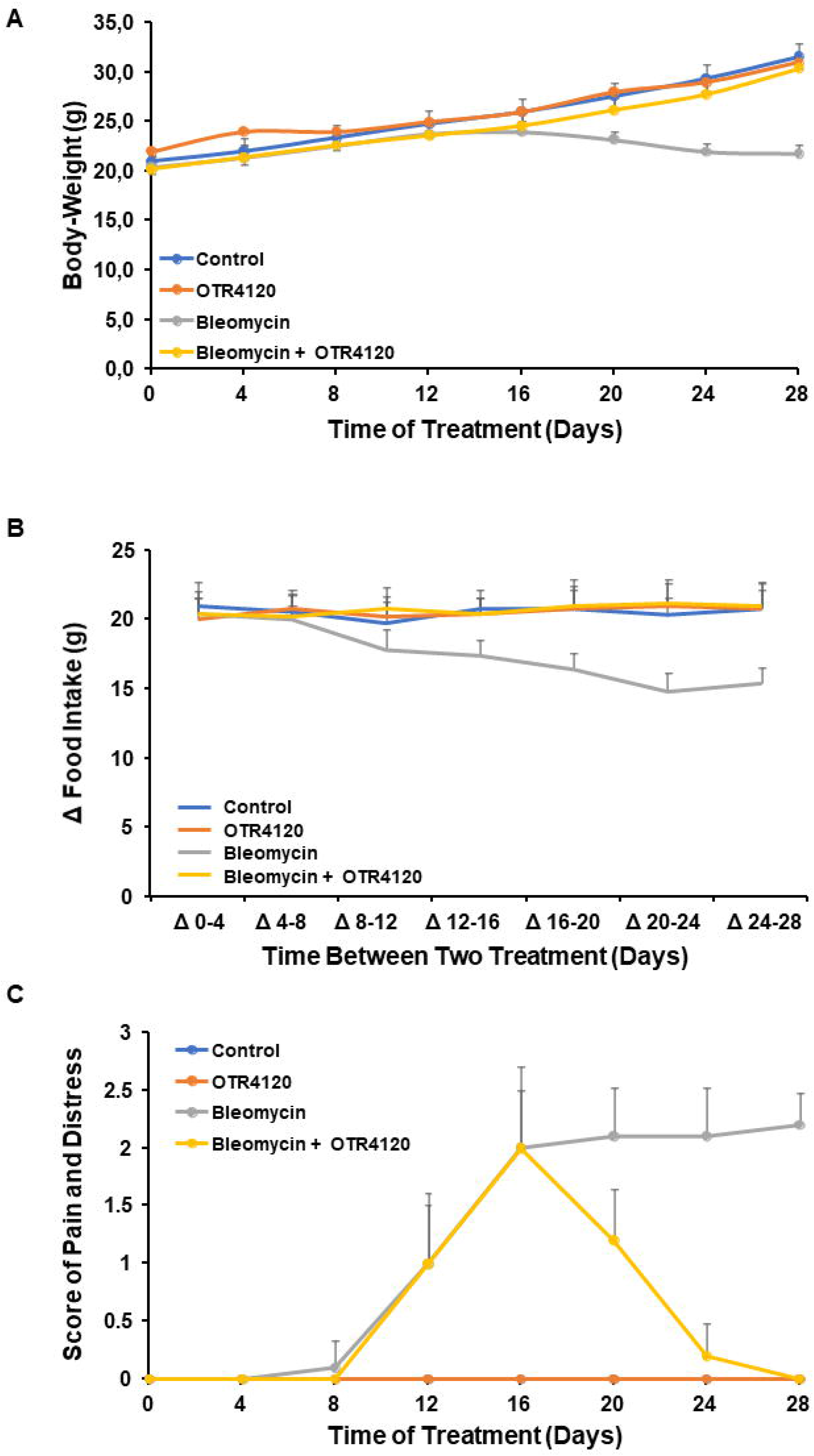
**Body weight (A), difference (Δ) in food intake (B), and score on pain and distresse (C)** of mice treated with bleomycin: Bleomycin (0.1 mg/kg) was injected I.P. at day 0 and 2. OTR4120 was injected at day 11, 14, 17, 21 and 24. Data presented as mean ± SD. Control (treated with saline only);OTR4120 only (1 mg/kg); Fibrosis model (bleomycin-induced pulmonary fibrosis); OTR4120-treated (bleomycin induced pulmonary fibrosis treated with OTR4120, 1 mg/kg). △*p-value*< 0.01 versus control group; \*\**p-value*< 0.01 versus fibrosis model group.

### OTR4120 reduced the increase in lung index scores induced by bleomycin

Lung/body weight index values in the fibrosis model group were significantly higher than those observed in the control group, at day 28 (9.97 ± 0.65 versus 7.31 ± 0.45 mg/g; fibrotic group versus controls, respectively; all *p-value*< 0.01; **Table 1**). In the OTR4120-treated fibrosis group, lung index values were significantly lower than in the fibrosis model group (**Table 1**). Lung index values were significantly lower in the OTR4120-treated group than in the fibrosis model group (7.63 ± 0.58 versus 9.97 ± 0.65 mg/g, *p-value*< 0.05], respectively; **Table 1**). There were no statistically significant differences between the control group and the OTR4120-treated group (7.31 ± 0.45 vs 7.63 ± 0.58 mg/g, *p-value*= non-significant).

**Table 1:**
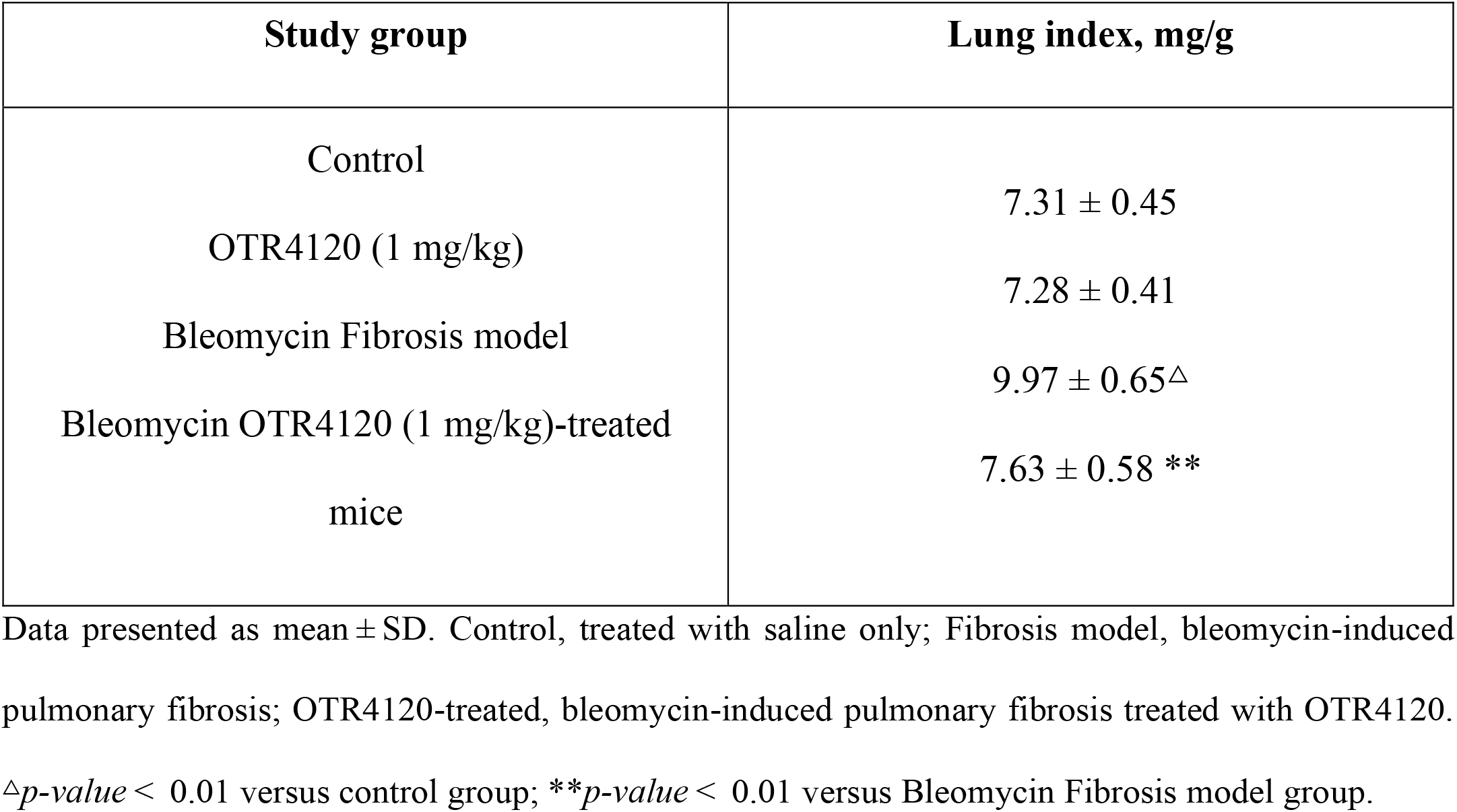
Lung index values in a bleomycin-induced Swiss mouse model of pulmonary fibrosis, treated with OTR4120.

| Study group | Lung index, mg/g |
| --- | --- |
| Control | 7.31 ± 0.45 |
| OTR4120 (1 mg/kg) | 7.28 ± 0.41 |
| Bleomycin Fibrosis model | 9.97 ± 0.65<sup>Δ</sup> |
| Bleomycin OTR4120 (1 mg/kg)-treated mice | 7.63 ± 0.58 ** |

Data presented as mean ± SD. Control, treated with saline only; Fibrosis model, bleomycin-induced pulmonary fibrosis; OTR4120-treated, bleomycin-induced pulmonary fibrosis treated with OTR4120.
<sup>Δ</sup>*p*-value < 0.01 versus control group; \*\**p*-value < 0.01 versus Bleomycin Fibrosis model group.

### OTR4120 treatment improves bleomycin-induced changes in lung histopathology

Haematoxylin and eosin-stained lung tissue sections were evaluated for histopathological abnormalities on days 28 following OTR4120 treatment (**Figure 3**). Under light microscopy, lung tissue sections from the control group showed normal alveolar spaces, normal thickening of the alveolar septa and high elasticity (Figure 3a and 3b). Lung sections from mice in the bleomycin-induced fibrosis group showed marked histopathological abnormalities, with alveolo-interstitial inflammation, at day 28 (**Figure 3c**). Lung sections from mice treated with OTR4120 showed moderate reduction of inflammatory cell infiltration, with normal alveolar structure and a few macrophages, lymphocytes and plasma cells (Figure 3d). Alveolitis/pulmonary fibrosis scores were significantly lower in the bleomycin-induced fibrosis model group versus the control group at days 28 (31.51 % vs 56.64 %; *p-value* < 0.01; **Table 2**). Decreased alveolitis/pulmonary fibrosis scores in response to bleomycin-induced fibrosis were significantly attenuated in mice treated OTR4120 (31.51 % vs 47.67 %; *P* < 0.05; Table 2).

**Table 2.** Alveolitis area scores (%).

| Study group | Alveolitis area scores (%) |
| --- | --- |
| Control | 56.64 ± 4.5 |
| OTR4120 | 55.94 ± 3.5 |
| Fibrosis model | 31.51 ± 3.2 $\Delta$ |
| OTR4120-treated bleomycin-induced fibrosis | 47.67 ± 5.6 ** |

**Figure 3.**
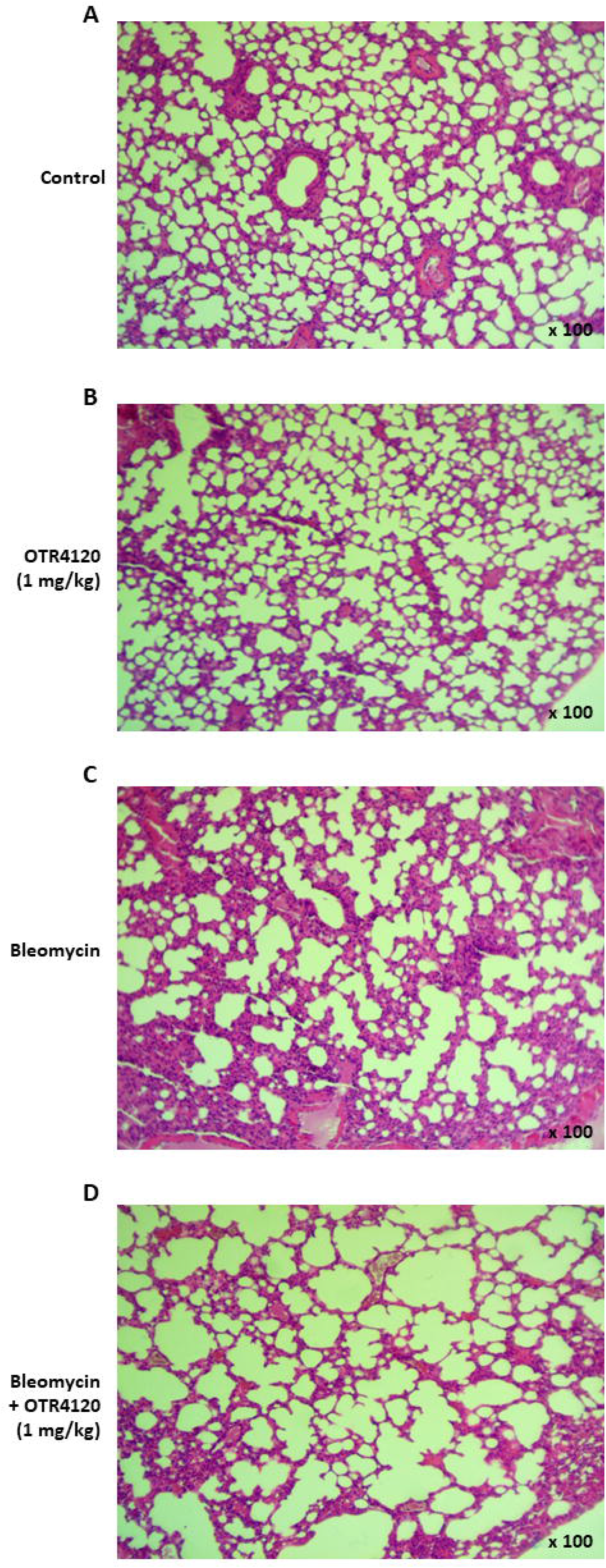
Representative photomicrographs showing haematoxylin and eosin (H&E) stained lung sections. Lung sections were obtained from a bleomycin-induced Swiss mouse model of pulmonary fibrosis: day 28 following initiation of OTR4120 treatment. Three study groups included (**A**)controls treated with saline only, (**B**)OTR4120 only (1mg/kg), (**C**)bleomycin-induced fibrosis model (0.1 mg/kg), and (**D**)bleomycin-induced fibrosis (0.1 mg/kg) treated with 1 mg/kg of OTR4120. Original magnification, x100.

Alveolitis area scores (%) in a bleomycin-induced Swiss mouse model of pulmonary fibrosis, treated with OTR4120 were assessed by Mesurim Software (http://acces.ens-lyon.fr/acces/logiciels/applications/mesurim). Data presented as mean ± SD. Control, treated with saline only; Control, treated only with OTR4120; fibrosis model, bleomycin-induced pulmonary fibrosis; bleomycin-induced pulmonary fibrosis treated with OTR4120. △*P* < 0.01 versus control group; \*\**P* < 0.01 versus fibrosis model group.

### OTR4120 reduced bleomycin-induced IL6 and TNF-α cytokine levels in the lung

To determine how OTR4120 affects the bleomycin-induced cytokine production, we measured the concentrations of TNF-α and IL-6 in BALF by ELISA. As illustrated in **Figure 4**, TNF-α (**Figure 4a**) and IL-6 (**Figure 4b**) levels were significantly higher in the experiment group than in the control group. OTR4120 (1 mg/kg) efficiently reduced the production of TNF-α and IL-6 compared to those obtained in the Bleomycin group (**Figure 4**).

**Figure 4.**
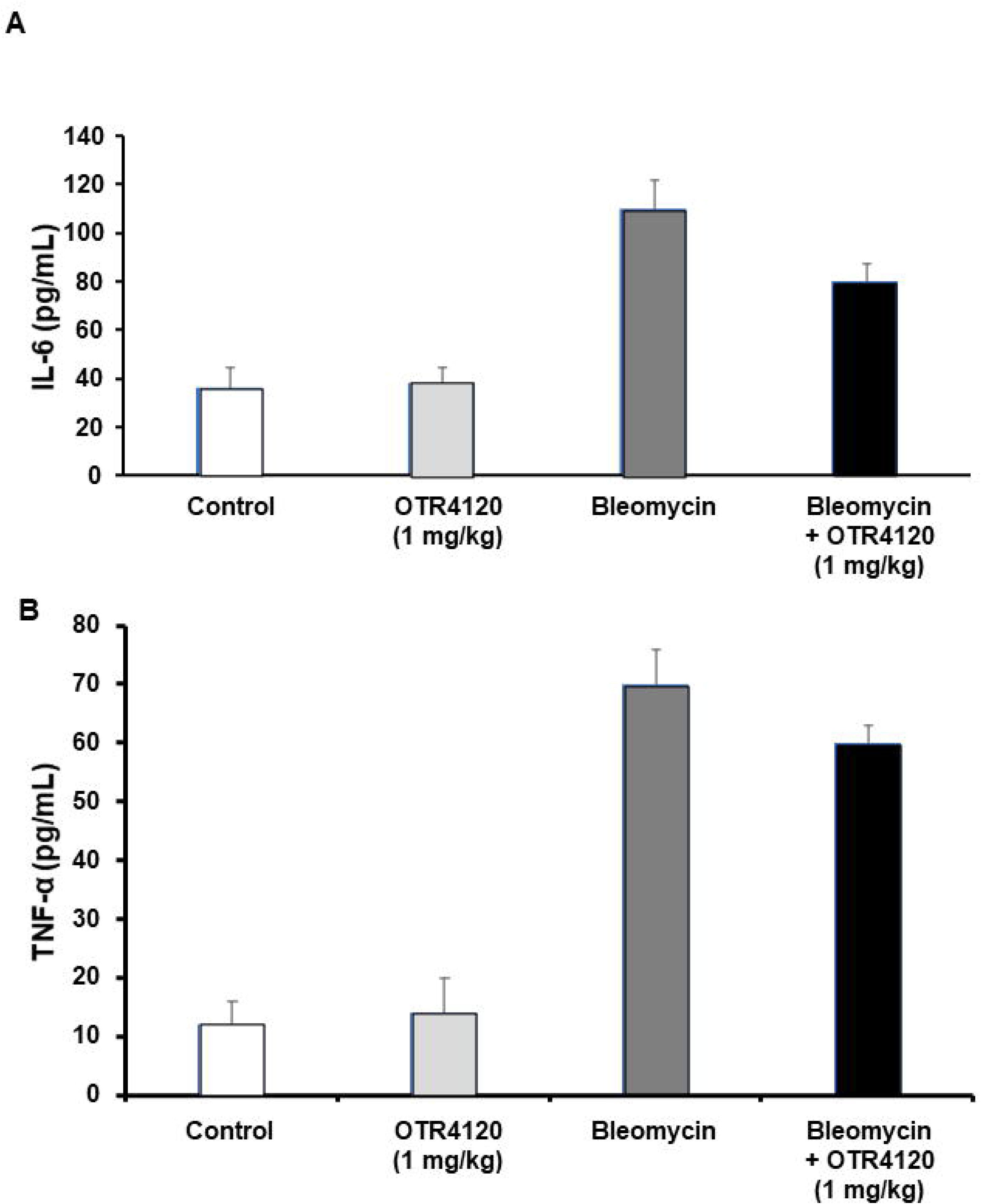
OTR4120 significantly attenuated the Bleomycin-induced (A) TNF-α and (B) IL-6 . Data presented as mean ± SD. * p-value<0.05 vs. Control group, # p-value<0.05 vs. Experiment group, p-value<0.05 vs. OTR4120 group. N=5.

### Effect of OTR4120 on the Collagen I and Collagen III levels in mice with bleomycin-induced pulmonary fibrosis

Collagen I and collagen III contents from these tissue specimens were quantitatively assessed by the Collagen I and Collagen III ELISA tests, and the production of each collagen type was determined. Collagen I production remained significantly unchanged when comparing control group to bleomycin-induced fibrosis group (48.00 ± 2.58 vs 50.00 ± 2.12 ng/ml) or with bleomycin-induced fibrosis group treated with OTR4120 (48.00 ± 2.58 vs 47.00 ± 3.12 ng/ml; **Figure 5a**).

**Figure 5.**
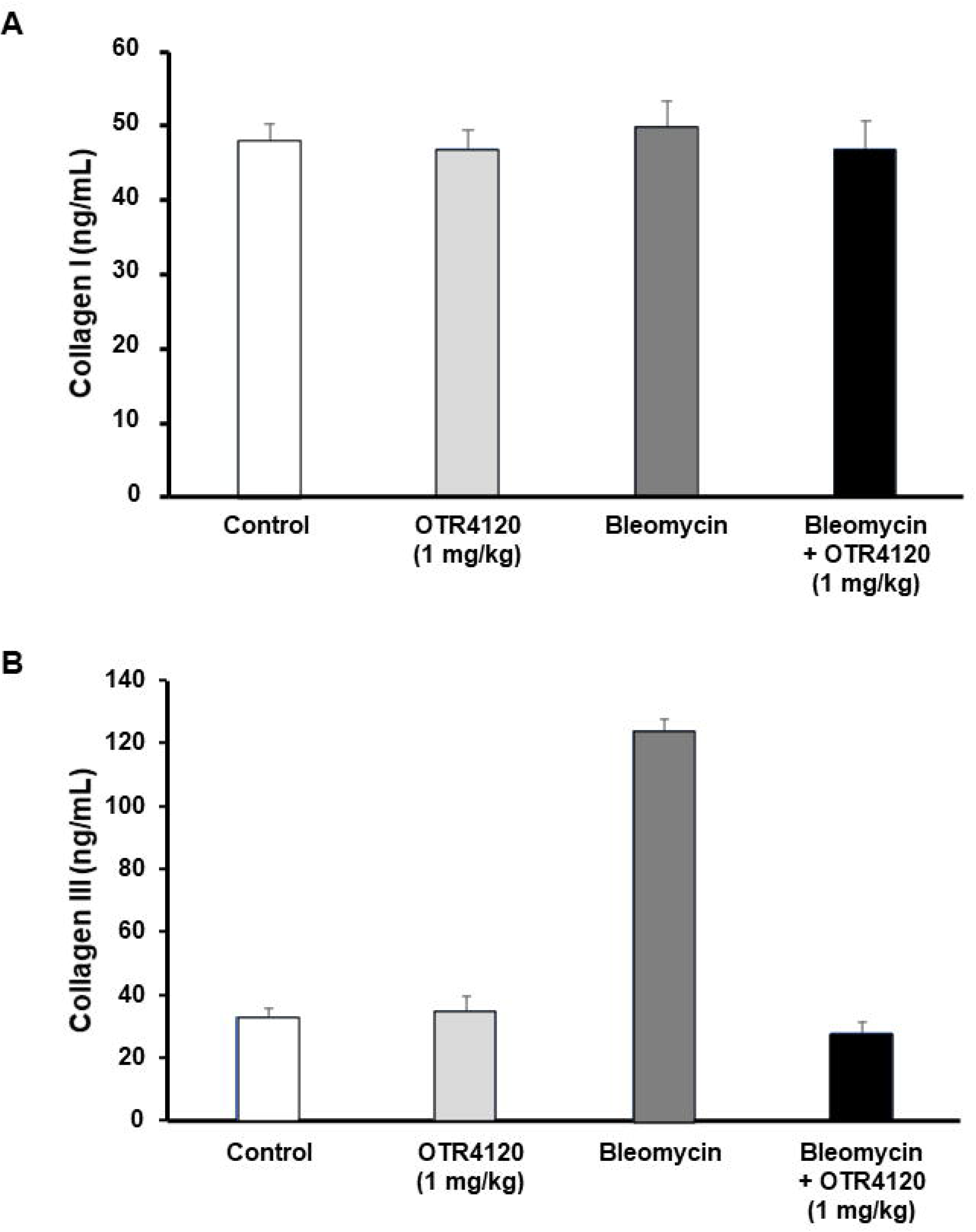
Quantification of soluble collagen. (**A**) Quantification of soluble collagen I in mice lung homogenates (Control, treated with saline only; OTR4120 only (1 mg/kg); bleomycin-induced pulmonary fibrosis (0.1 mg/kg); bleomycin-induced pulmonary fibrosis (0.1 mg/kg) treated with OTR4120 (1 mg/kg). **(B**) Quantification of soluble collagen III in mice lung homogenates (Control, treated with saline only; OTR4120 only, bleomycin-induced

In contrast, collagen III levels were found to be significantly increased in lungs from the bleomycin-induced fibrosis group compared with the control group, (124.28 ± 1.37 versus 33.00 ± 1.79 ng/ml; *p-value* < 0.05; **Figure 5b**). Treatment with OTR4120 was found to significantly attenuate the increased Collagen III levels associated with bleomycin administration (124.28 ± 1.37 versus 28.00 ± 2.10; *p-value* < 0.01; **Figure 5b**). There were no statistically significant differences in lung Collagen III levels between the control group and the bleomycin-induced fibrosis group treated with OTR4120 (33.00 ± 1.79 versus 28.00 ± 2.78 ng/ml; *p-value*= non-significant; **Figure 5b**).

## Discussion

Pulmonary fibrosis in clinics can result from various causes that may not always be well identified; some causes may be repetitive microinjury of the alveolar epithelium, hypoxia, viral or microorganisms destruction, inefficient and/or aberrant epithelial repair, excessive extracellular matrix deposition and destruction of distal lung architecture, etc. In all cases, the cellular micro-environment is destroyed and the tissue responses are often characterized by increased number of polymorphonuclear cells (PMNs), alveolar and interstitial cell infiltrates in the lungs, and the presence of edema [26], consistent with features of the bleomycin-induced pulmonary fibrosis model in the present study. Thus, in line with the frequent use in research, we show here that bleomycin-induced pulmonary fibrosis can be used to explore new pharmacological methods to treat reduce pulmonary fibrosis [27–28]. Indeed a common criticism in the bleomycin-induced pulmonary fibrosis model is the role of inflammation, as it induces chronic inflammation and fibrosis, evidenced by the release of various inflammatory mediators [29–30]. In the present study, bleomycin treatment was also found to cause evident chronic inflammation and fibrosis compared with the saline-treated control group, and this increased inflammation and fibrosis was significantly decreased by OTR4120 treatment. H&E-stained lung tissue sections from the control group showed that normal alveolar spaces, and normal thickening of the alveolar septa were present. However, sections from the lungs of mice in the bleomycin-induced fibrosis group showed marked histopathological abnormalities. Tissue sections from mice treated with OTR4120 showed amelioration of inflammatory cell infiltration, together with a reduction in interstitial thickening.

The majority of animal models of fibrosis including the above-described model starts with inflammation [31]. TNF-α is a key cytokine in pulmonary fibrosis, which induces expression of adhesion molecule, recruitment of inflammatory cells into the lungs and synthesis of other cytokines [32–33]. More recently Jia-Ni Zou [34] demonstrated significant relationships between pulmonary fibrosis and the levels of albumin and IL-6, which suggest that IL-6 and albumin are independent risk factors affecting pulmonary fibrosis. Previous studies reported that IL-6 could serve as an indicator of the progression of COVID-19 [35]. IL-6 levels can reflect the severity of the inflammatory response. In our study, BLM-induced model is characterized by increased levels of TNF and IL-6 in homogenated lungs. OTR4120 significantly reduced the increases in TNF-α and IL-6 levels in both models. Therefore, these results suggest that OTR4120 might provide the protection from lung fibrosis by an anti-inflammatory effect. It is generally accepted that the regulation of pro-inflammatory cytokines is mediated, at least partly, by NF-κB and that the abnormal humoral metabolism and pulmonary edema contribute to the severity of symptoms and fatality of COVID-19 patients. Decreased expression of alveolar Na-K-ATPase, dysregulation of sodium and potassium channels, aquaporins, and renin angiotensin system, and abnormal metabolism of bradykinin and hyaluronic acid as well as cytokine inflammatory storm all lead to ARDS and pulmonary edema [36].

Cytokines activate neutrophils and promote their accumulation at the site of injury, induce the release of protease and oxygen free radicals, and finally lead to pulmonary interstitial edema and a severe inflammatory response [26,37]. To the best of the authors’ knowledge, the present study is the first to show the effect of OTR4120 on lung index values in mice. Lung index values in the bleomycin-induced fibrosis group were increased, and the magnitude was significantly reduced in the OTR4120 treated group, suggesting that OTR4120 attenuates bleomycin-induced lung injury in mice.

Many studies reported that increased type III collagen serum levels are representative of active fibrosis [38,39]. An increase in type III collagen was observed in lung biopsy specimens taken from patients in the early stages of cryptogenic pulmonary fibrosis [39] and from subjects with active fibrotic disease. Shahzeidi, et al. [40] suggest that type III procollagen gene expression was enhanced in bleomycin induced fibrosis and that expression was maximal between 10 and 35 days after a single dose of bleomycin. The most active cells were located in interstitial areas around the conducting airways, although these cells were usually seen in areas with no histological evidence of fibrosis. Regions with the most advanced fibrosis, as assessed by histological methods, rarely contained cells with activity above the threshold detectable by this technique. These results suggest that activation of interstitial fibroblasts, with enhanced type III collagen gene expression, forms at least part of the mechanism leading to increased collagen deposition in bleomycin induced fibrosis and that this occurs before fibrosis is detected by conventional histological staining.

Our results demonstrated that type-III collagen levels were significantly increased in BLM treated lungs compared to those in saline-treated or BLM-untreated control lungs. The enhanced type-III collagen levels in BLM-treated lungs can be obviously ameliorated by OTR4120 treatment, suggesting an antifibrotic effect of OTR4120. This specific effect of OTR4120 on collagen III synthesis has been previously described [41].Antifibrotic effects of OTR4120 have been demonstrated in earlier studies [42]. RGTA treatment has been shown to decreased the production of collagen III in intestinal tissue specimens from patients with Crohn’s disease [43]. In the present novel investigation of a bleomycin-induced pulmonary fibrosis mouse model that compared OTR4120-treated mice versus untreated mice with bleomycin-induced fibrosis, we demonstrated that OTR4120 exerts an anti-inflammatory effect in the lungs of mice with bleomycin-induced pulmonary fibrosis. These effects were evidenced by attenuation of the morphological fibrotic responses, and the reduced lung index values, neutrophil infiltration and collagen levels.

During the restoration period, the clinical signs induced by bleomycin administration in most animals resolved gradually 17 days after Bleomycin administration and 6 days after OTR4120 treatment and disappeared almost completely at days 24.

Further studies are needed to confirlm the results and clarify the mechanism with which OTR4120 attenuates bleomycin-induced pulmonary fibrosis

## Conclusion

The present research revealed that OTR4120 could efficiently attenuate bleomycin-induced pulmonary fibrosis in a mouse model. The study provides evidence that OTR4120 may be a promising candidate to treat patients suffering from lung injuries.

This concept was supported in a recenr case report on13 patients with Covid and lung lesions treated by nebulized OTR4120 [44]. It provided the first human safety case data of OTR4120 by airway administration with promising trends of clincal improvements and paved the way to placebo controlled trials.

